# The first RNA viruses detected in a trypanosome: novel narnaviruses and a leishmaniavirus in the gecko parasite *Trypanosoma platydactyli*

**DOI:** 10.64898/2025.12.11.693634

**Authors:** Alexei Yu. Kostygov, Danyil Grybchuk, Donnamae Klocek, Jan Votýpka, Jairo A. Mendoza-Roldan, Julius Lukeš, Vyacheslav Yurchenko

## Abstract

**Background:** Trypanosomatids are parasitic flagellates best known for human pathogens causing sleeping sickness, Chagas disease, and leishmaniasis. RNA viruses infecting these protists have recently gained attention for their role in disease severity. While numerous such viruses have been described in *Leishmania* and several other trypanosomatid genera, none have previously been documented in the iconic genus *Trypanosoma*.

**Results:** We report the first discovery and molecular characterization of RNA viruses in trypanosomes, identifying a leishmaniavirus and two narnaviruses in a single strain of *Trypanosoma platydactyli*, a parasite of the common wall gecko. The leishmaniavirus genome revealed a conserved organization, including a putative ribosomal frameshift site and a hairpin-like secondary structure typical of the genus. Phylogenetic inference indicates that it is closely related to leishmaniaviruses from Old World *Leishmania* spp., consistent with shared vector ecology. The two narnaviruses have distinct origins, although both cluster with viruses of other trypanosomatids, suggesting historical exchanges among co-infecting parasites.

**Conclusions:** Our study expands both the known diversity of RNA viruses in trypanosomatids and the range of trypanosomatid genera that host these viruses, providing guidance for future screening. We suggest that vector ecology—particularly feeding behavior—may influence viral acquisition by trypanosomes, explaining the previous absence of viral reports from intensively studied trypanosomes of medical relevance vectored by tsetse flies or kissing bugs. Therefore, overlooked species transmitted by Nematocera represent promising candidates for future viral discovery. This concept extends beyond trypanosomatids, providing a general framework for understanding the conditions that permit viral host switching by viruses among microeukaryotes.

## Background

Flagellates of the family Trypanosomatidae represent one of the best-studied groups of protists, known primarily for human pathogens, such as *Trypanosoma brucei*, *T. cruzi*, and several species of the genus *Leishmania*, which cause sleeping sickness, Chagas disease, and various forms of leishmaniasis, respectively [1]. Trypanosomes, leishmaniae, and members of the phytoparasitic genus *Phytomonas* are dixenous parasites, which require an invertebrate (typically, insect) vector for transmission, whereas the majority of trypanosomatid genera are monoxenous, developing exclusively in insects [2].

Due to the strong interest in trypanosomatids, they were among the first protists screened for RNA viruses using molecular methods [3]. The first such viruses were discovered in *Leishmania* (*Viannia*) *guyanensis* [4, 5] and in a related species – *L.* (*V.*) *braziliensis* [6]. These double-stranded RNA viruses were classified into the newly established genus *Leishmaniavirus* originally placed in the family *Totiviridae* [7] and now assigned to *Pseudototiviridae* [8]. Shortly thereafter, another member of this genus was discovered in *Leishmania major* [9]. Additionally, virus-like particles resembling RNA virions have been detected in various trypanosomatids using electron microscopy [3], although most of these observations remain unconfirmed at the molecular level.

Interest in trypanosomatid-infecting viruses has grown significantly following the discovery that the presence of leishmaniaviruses (LRVs) causes elevated expression of proinflammatory cytokines and aggravates the course of leishmaniasis, making it metastatic [10–12]. This prompted targeted screening of other trypanosomatids, which resulted in identification of many novel viruses across several lineages. These include *Mundinia* and *Sauroleishmania*, two neglected subgenera of *Leishmania* [13, 14]; other Leishmaniinae [15–20], as well as more distantly related genera *Blechomonas*, *Phytomonas*, and *Obscuromonas* [21–23].

Within the last decade, there has been a surge in the number of known viral species inhabiting trypanosomatids, rising to over thirty from the original two (or three, as LRV in *L. aethiopica* was recently established as a separate species [8]). In addition to novel leishmaniaviruses, which proved to inhabit not only *Leishmania* spp. but also the phylogenetically distant monoxenous genus *Blechomonas* [21], other viral groups were discovered in trypanosomatids. These include positive-sense single-strand RNA (ssRNA) viruses from the families *Narnaviridae*, *Mitoviridae*, and a group of tombus-like viruses; negative-sense ssRNA viruses from *Qinviridae* and *Leishbuviridae*; as well as the enigmatic, unclassified genus *Ostravirus* [15, 17, 23, 24]. The monoxenous species *Leptomonas pyrrhocoris* (subfamily Leishmaniinae) has proven especially prone to viral infections, with prevalence reaching 60%, seven viral species from four family-level taxa recorded, and up to four different viruses co-occurring in a single isolate.

Despite the growing number of RNA viruses discovered in various trypanosomatid lineages, none have been molecularly characterized in any members of the genus *Trypanosoma*. To date, there were only two electron-microscopy based reports of virus-like particles in *Trypanosoma melophagium* and *T. cruzi* [25, 26]. However, the viral nature of these particles remains unconfirmed. In the case of *T. melophagium*, only a single electron-dense structure—interpreted as a potential virion—was observed, which could also constitute an artifact.

Here, we present the first discovery and molecular characterization of RNA viruses in the common wall gecko parasite *Trypanosoma platydactyli*. Based on our findings, we outline potential reasons why RNA viruses are rare in other trypanosomes and discuss the ecological context of virus acquisition in trypanosomatids.

## Results

### Morphology of the studied trypanosome strain

Although this study focuses on viruses, confirming the host species identity was essential. Prior to the isolation of strain RI-340, no reference molecular data were available for *Trypanosoma platydactyli* to enable its easy and reliable identification. In a previous study, the genome of this strain was published [27], but no evidence was provided to support its species classification. Here, we resolve this issue by presenting data on the morphology of the studied trypanosome in the blood and compare it to the original description of *T. platydactyli*. Additionally, we provide a brief overview of morphotypes observed in culture.

Consistent with the original description [28], the trypomastigotes observed in the bloodstream of the lizard host were elongated, broadened in the posterior half, and markedly curved (Fig. 1A). They harbored a well-developed undulating membrane with several large folds, a short (approximately half the cell length) free flagellum, and closely positioned nucleus and kinetoplast. Cell dimensions generally matched those reported or illustrated in the original description (Additional file 1: Table S1), except for two parameters measured on Catouillard’s figure, which likely underrepresented natural variation due to a small sample size. Given the absence of other trypanosome species described from *Tarentola mauritanica* and the strong morphological similarity to the original description, the RI-340 strain can be unambiguously identified as *Trypanosoma platydactyli*.

**Fig. 1.**
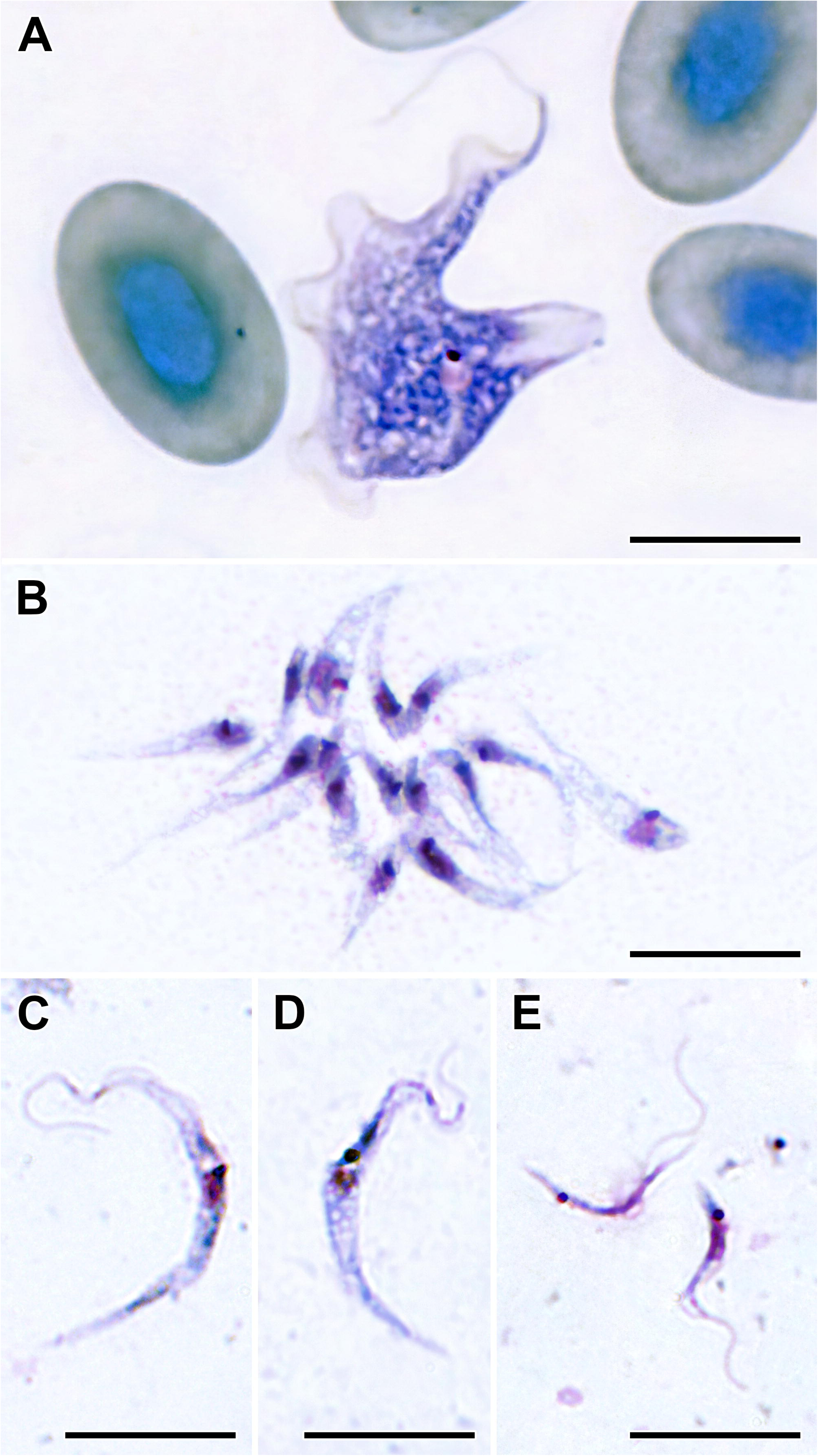
Giemsa-stained cells of the strain RI-340. (A) Trypomastigote in a blood smear. (B–E) Forms in culture: (B) rosette of tadpole-like cells; (C, D) epimastigotes; (E) trypomastigotes. Scale bars: 10 μm.

In culture, the cells were conspicuously smaller than in the bloodstream, particularly in width (Fig. 1B–E; Additional file 1: Table S2). Most abundant were tadpole-like cells without a visible flagellum, typically arranged in rosettes, with the kinetoplast and nucleus positioned close to the anterior end (Fig. 1B). In addition, two typical trypanosome morphotypes were observed: i) long, usually sickle-shaped epimastigotes with the kinetoplast adjacent to the nucleus near the cell center (Fig. 1C, D), and ii) very thin curvy trypomastigotes with variable positions of both DNA-containing structures (Fig. 1E).

Transmission electron microscopy revealed that, in addition to the typical trypanosomatid cell components (nucleus, kinetoplast, glycosomes, etc.), the RI-340 strain featured electron-lucent membranous vesicles—typically one per section—situated near the dictyosomes of the Golgi complex (Fig. 2A). These vesicles had slightly elliptical profiles (205–405 × 182–392 nm) and enclosed dark, rounded bodies approximately 30–55 nm in diameter (Fig. 2B), consistent with the known sizes of LRV virions [29–31]. Occasionally, these bodies were surrounded by a ring, presumably corresponding to a capsid shell (Fig. 2, inset).

**Fig. 2.**
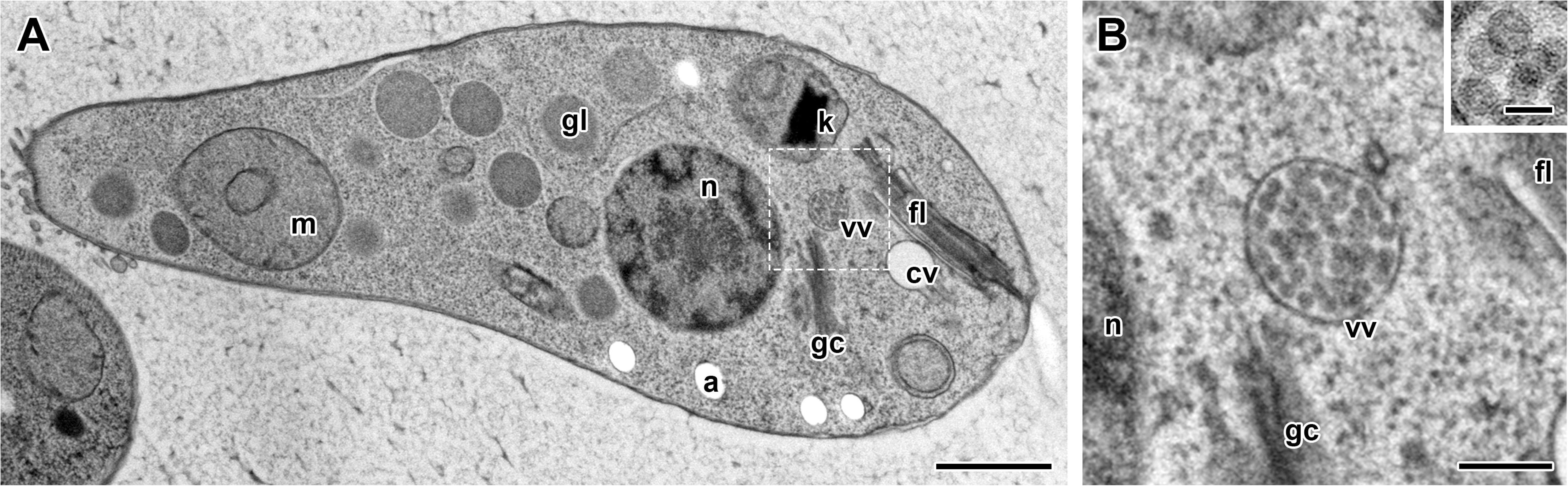
Transmission electron microscopy of the strain RI-340. (A) General view of a tadpole-like cell. The enlarged image of the boxed area is shown in (B). Inset: virus-like particles at higher magnification, showing an outer shell. Abbreviations: a – acidocalcisome, cv-contractile vacuole, fl – flagellum, gc – Golgi complex, gl – glycosome, k-kinetoplast, m – mitochondrion, n – nucleus, vv – vesicle with virus-like particles. Scale bars: (A) – 1 μm, (B) – 200 nm, inset – 50 nm.

### Viral screening and characterization

Electrophoresis of the double-stranded RNA (dsRNA) preparation from the RI-340 strain revealed three fragments: two approximately aligned with the 6 kb and 4 kb bands of the DNA ladder, and the third positioned slightly above the 3 kb band (Fig. 3A, Additional file 2: Fig. S1). The slower mobility of dsRNA in agarose gel compared to DNA of the same length explains the discrepancy with the size estimates from NGS data: 5.3, 3.7, and 2.8 kb (Fig. 3B; Additional file 1: Table S3). All three segments corresponded to distinct viral species—one leishmaniavirus and two narnaviruses—designated here as TplaLRV1, TplaNV1, and TplaNV2, respectively. The structure of their genomes was typical of the corresponding taxa. In both narnaviruses, most of the genome was occupied by a single RdRp-encoding ORF, with flanking regions of only 36–84 nucleotides in length. In TplaLRV1, three genes were predicted: a short ORF1 encoding a hypothetical protein and two large ORFs that overlap *via* a −1 ribosomal frameshift, enabling the production of either a capsid protein alone or a capsid–RdRp fusion protein [32]. A BLASTn search using ORF1 as a query returned hits in LRVs from *L. major*, *L. aethiopica*, and *L.* (*Sauroleishmania*) spp., with 91–93% coverage and 84.7–87.2% identity, indicating that this nucleotide sequence is highly conserved.

**Fig. 3.**
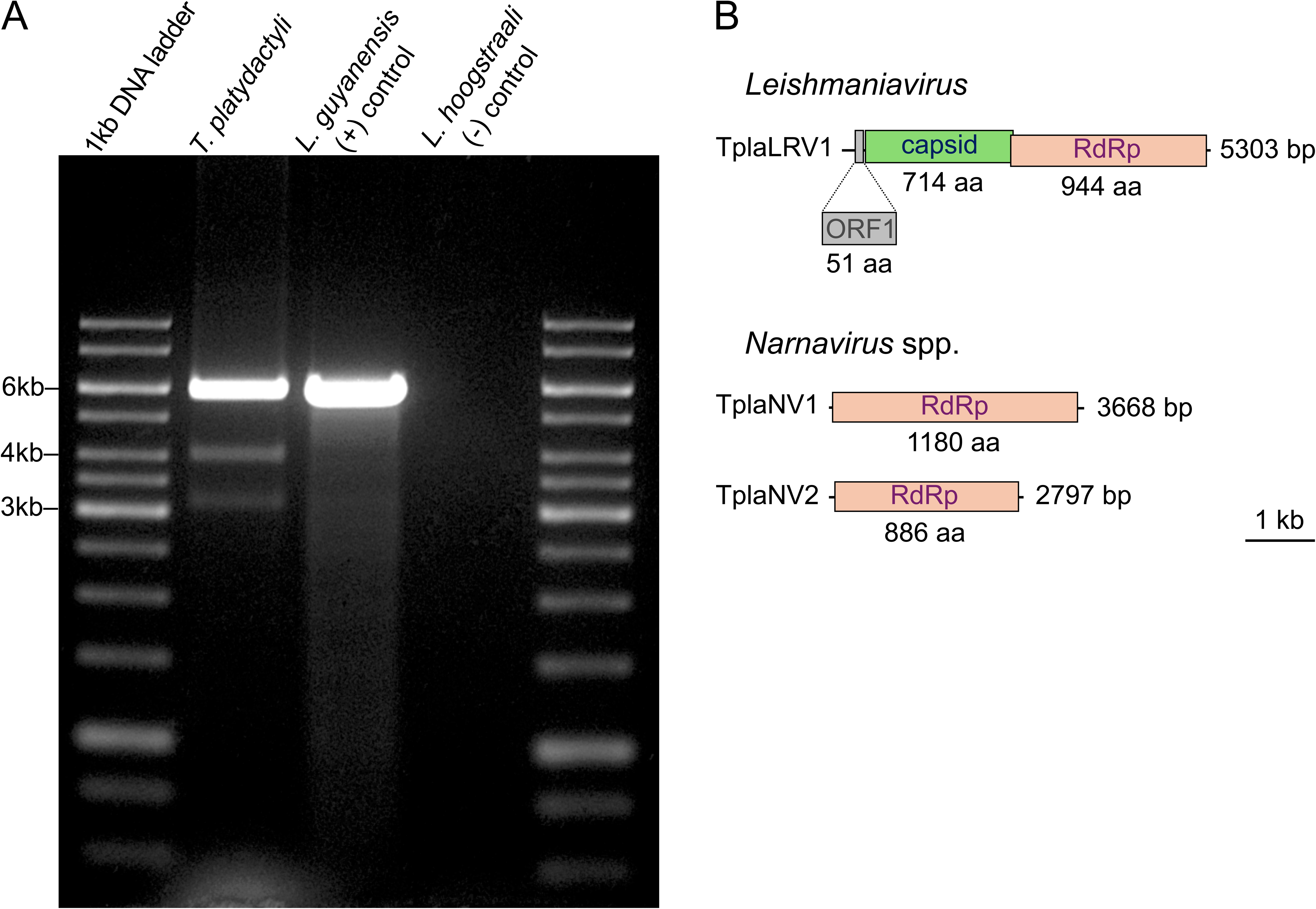
Viruses documented in the strain RI-340. (A) Screening for dsRNA. (B) Schematic representation of viral genomes, drawn to scale.

The exact position of the frameshift in the TplaLRV1 genome could not be determined. We applied the recently developed software PRFect [33], but it failed to identify any sequence associated with programmed ribosomal frameshift. Therefore, we delineated the region between the stop codon in the capsid frame and the nearest upstream stop codon in the RdRp reading frame (positions 2418–2469; Additional file 3: Suppl. Fig. S2) and analyzed it further. Within this region, IPknot++ predicted a hairpin (positions 2,428–2,460), a typical feature associated with the frameshift region in other LRVs [3]. Thus, the RdRp portion of the fused protein most likely starts between the positions 2,418 and 2,427. Previously, the identification of the ribosomal frameshift in the genome of LRV from *Leishmania aethiopica* (now classified as *Leishmaniavirus sani*) was not accompanied by any indication of whether the capsid-RdRp overlap region contained a hairpin [12], leaving the impression that it was absent [3]. To clarify if this secondary structure is also present in that LRV species, we applied the same approach and identified a hairpin separated from the upstream stop codon in the RdRp frame by only five nucleotides (Additional file 2: Fig. S1). Hence, it appears that this feature is universally present in LRVs.

### Phylogeny of leishmaniaviruses

In line with our previous inferences [14, 21], the phylogenetic reconstruction performed here strongly supported non-monophyly of leishmaniaviruses from *Leishmania*. They formed two distinct clades intermingled with viruses from *Blechomonas*: one associated with Old World species (*L. major*, *L. aethiopica*, and *L.* (*Sauroleishmania*) spp.) and another – with New World species of the subgenus *Viannia* (Fig. 4). The LRV from *Trypanosoma platydactyli* was recovered as sister to the first group, which likely reflects their shared biogeographical origin in the Old World and reliance on the same vectors – sandflies of the genera *Phlebotomus* and/or *Sergentomyia* [34–38].

**Fig. 4.**
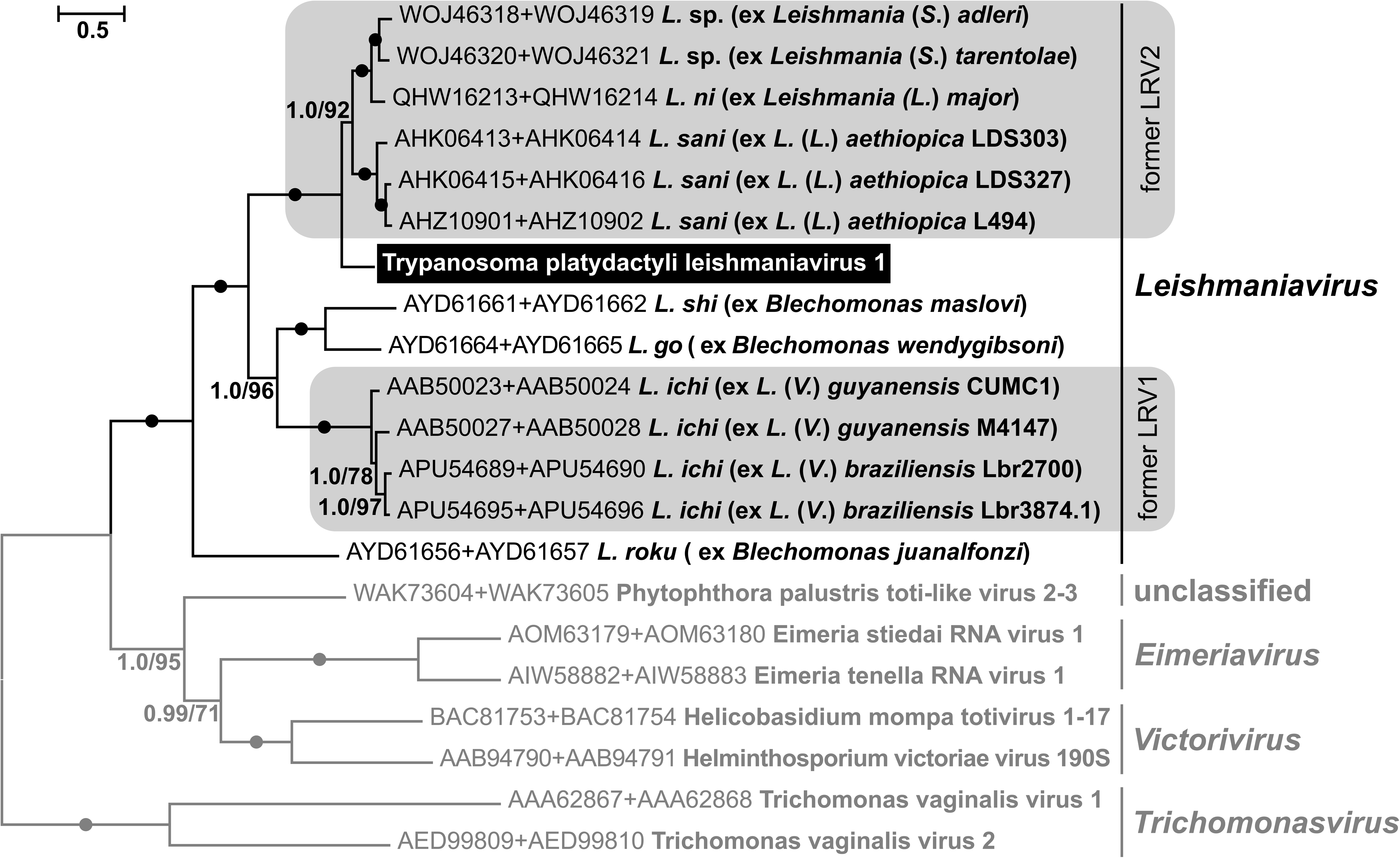
Maximum likelihood phylogenetic tree of the family *Pseudototiviridae* based on concatenated amino acid sequences of the capsid and RdRp proteins. The tree is rooted according to a previous inference [14]. Taxa outside *Leishmaniavirus* are shown in grey; the LRV reported in this study is highlighted in black. Numbers at branches represent Bayesian posterior probabilities and bootstrap supports; black circles indicate full support (1.0/100).

Considering frequent coinfections of *T. platydactyli* and *L. tarentolae*, which in the past even led to confusion over whether these two species are distinct [39, 40], one would expect their LRVs to be closely related. However, the strongly supported phylogenetic position of the virus from *T. platydactyli* indicates that it has a more ancient origin than those infecting *Sauroleishmania* (Fig. 4). This suggests that *T. platydactyli* is not the only LRV-bearing trypanosome and that related viruses may infect other species of the subgenus *Squamatrypanum*, to which it belongs [2, 41].

### Phylogeny of narnaviruses

Our inference of *Narnaviridae* phylogeny revealed two strongly supported clades, comprising viruses from trypanosomatids, as well as those from a diverse array of plants, animals, fungi, and other hosts (Fig. 5). The two narnaviruses from *T. platydactyli* were found to have distinct origins, each falling within one of the two clades. However, in both cases, they unambiguously grouped with viruses from other trypanosomatids.

**Fig. 5.**
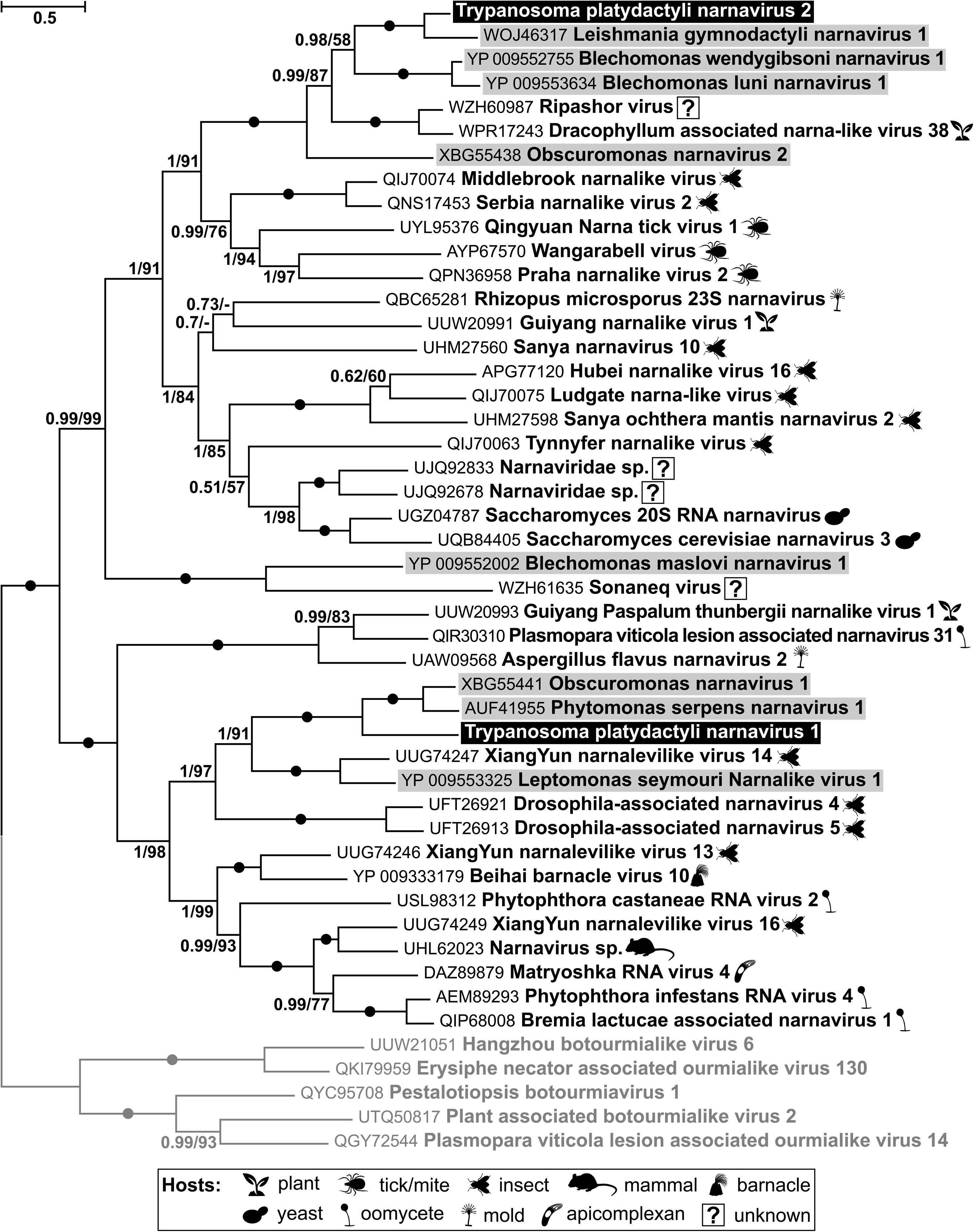
Maximum likelihood phylogenetic tree of *Narnaviridae* based on RdRp amino acid sequences. The tree is rooted with *Botourmiaviridae* (shown in grey). The two viruses reported in this study are highlighted in black, while other narnaviruses from trypanosomatids are highlighted in grey. For the others, isolation sources (presumed hosts) are shown with pictograms explained in the graphical legend. Numbers at branches represent bootstrap supports and Bayesian posterior probabilities; circles indicate absolute support (100/1); bootstrap values below 50% are replaced with dashes.

TplaNV1 was positioned as sister to viruses from *Obscuromonas* sp. and *Phytomonas serpens*, while also showing a close relationship to that from *Leptomonas seymouri* (Fig. 5). This suggests that *T. platydactyli* acquired this narnavirus from other trypanosomatids, although the exact donor cannot be presently identified—especially given that *Phytomonas* and *Obscuromonas* do not inhabit sandflies. Instead, they parasitize true bugs, and (in the case of the former) also plants. Other viruses branching nearby have been detected in mosquitoes and fruit flies [42, 43], suggesting a possible origin of this group from insect-associated narnaviruses. However, these viruses may be hosted by trypanosomatids, as the NCBI Taxonomy Analysis reports for the corresponding metatranscriptomic SRAs indicate the presence of sequences from these flagellates [44–49].

TplaNV2 nested within a clade of trypanosomatid-associated narnaviruses (one from *Leishmania* and two from *Blechomonas* spp.), and, notably, was positioned as sister to the virus from *L.* (*Sauroleishmania*) *gymnodactyli.* This suggests that, unlike LRVs, narnaviruses have been directly exchanged between sandfly-and lizard-infecting sauroleishmaniae and trypanosomes, an event that apparently occurred recently in evolutionary terms. The closest relatives of the entire trypanosomatid-infecting clade comprising TplaLRV2 are two viruses reported from a plant and an environmental sample [50, 51]; however, the corresponding SRAs contained trypanosomatid signatures as evidenced in NCBI Taxonomy Analysis reports, as above [52, 53]. Given this and the fact that the next closest neighbor on the phylogenetic tree infects the trypanosomatid *Obscuromonas* sp., it is parsimonious to suggest that the clade encompassing all seven viruses discussed here is associated with trypanosomatids. Its sister clade comprises viruses exclusively found in arthropods, suggesting that, in this part of the tree, trypanosomatids acquired narnaviruses from insects. Ticks and mites are less likely donors, as they are not common hosts/vectors for trypanosomatids and are restricted to just a few species of these parasites [2].

## Discussion

Given that no viruses have previously been characterized in the genus *Trypanosoma* that unites over 500 described species [2], our discovery of three viruses at once in a single strain of *T. platydactyli* is particularly interesting and unexpected. What might explain this long-standing gap in knowledge? One key reason is that only a limited number of *Trypanosoma* spp. were introduced into culture and investigated in more detail upon their description. Research has primarily focused on a few species of practical importance, such as human pathogens *T. cruzi* and *T. brucei*, nonpathogenic *T. rangeli*, as well as salivarian trypanosomes responsible for livestock diseases, including *T. congolense*, *T. vivax*, and others [54]. Why, then, have no viruses been identified at least in these intensively studied trypanosomes?

To answer this question, one should consider the possible viral donors and the conditions necessary for viral transition. By “transition” we mean the evolutionary switch from one host species to another, which requires subsequent adaptation and ultimately the establishment of a new viral species. To increase the likelihood of such a transition, the donor and recipient should frequently co-occur in close proximity. Infections must happen within a host or a vector of the flagellates, as trypanosomes generally do not appear in external environments or, if discharged there, survive only briefly [55, 56]. Potential donors include the host or vector, as well as their endosymbionts (including other trypanosomatids) and, in the case of the latter, ingested food. To the best of our knowledge, there is no evidence that parasitic protists (including trypanosomatids) have ever acquired viruses from vertebrates. For RNA viruses this would likely be too great an evolutionary leap, not achievable through stepwise adaptation, although such a possibility cannot be entirely excluded. Moreover, vertebrates are also unlikely to serve as sites for viral exchange, as their large body size makes them too loose an environment for the close interactions required for the potential viral transfer. In contrast, insects, acting as vectors for dixenous and hosts for monoxenous trypanosomatids, offer a much more favorable setting. Within its relatively small intestine, trypanosomatids often develop in very large numbers [57]. Thus, they can come into contact with other resident inhabitants of the gut and transient material passing through as food (usually of plant or animal origin) or as its microbial contaminants, such as yeasts in sap or oomycetes in decaying plant tissue. Hence, in the gut, numerous trypanosomatid cells can be efficiently exposed to viruses from various sources, but how much this potential is realized largely depends on the feeding habits of the insect host/vector. The most remarkable example in this respect is *Leptomonas pyrrhocoris*, which inhabits the omnivorous firebug *Pyrrhocoris apterus* and leads both in the overall number of described viruses and those detected simultaneously [17].

As for trypanosomes, the vectors of the well-studied species and *T. platydactyli* are fundamentally different. The American species (*T. cruzi* and *T. rangeli*) are transmitted by kissing bugs (Triatominae), whereas the African (salivarian) species are vectored by tsetse flies (*Glossina* spp.) [54]. Although sandflies are formally classified as hematophagous, this applies only to females during egg production, whereas for most of their lives, both sexes rely on plant sugars, such as sap, nectar, and honeydew [58]. This creates favorable conditions for infection by monoxenous trypanosomatids, as these food sources are frequently contaminated with feces from other insects [57]. Sandfly larvae are detritivorous [59] and may theoretically become infected with trypanosomatids, although such cases have not been reported. In contrast, tsetse flies are considered strictly hematophagous, with both sexes exhibiting this habit throughout their adult life, while the larva develops within the mother’s body and is deposited shortly before pupation [60, 61]. Although starved tsetse flies have been shown to consume small amounts of sugar under laboratory conditions [62], this is unlikely to be ecologically significant. The activity of their carbohydrate-catabolizing enzymes is markedly reduced compared to other dipterans, and energy metabolism relies primarily on amino acids, especially proline [61].

Notably, some salivarian trypanosomes have alternative transmission routes that further reduce the likelihood of viral acquisition. These include: (i) mechanical transmission by tabanids and *Stomoxys* flies—the sole route for *T. brucei evansi* and an alternative to biological transmission by tsetse for *T. vivax* and *T. congolense* [63–67]; as well as (ii) direct sexual transmission in *T. brucei equiperdum* [68].

In hematophagous triatomine bugs (also known as kissing bugs), males, females, and nymphs generally feed on blood, which is essential for the key life cycle events, such as molting and egg production [69]. However, exceptions exist, particularly in species that have not yet fully transitioned from predation to hematophagy [70]. These include: i) cannibalism; ii) cleptohematophagy, i.e., feeding on blood from engorged bugs; iii) haemolymphagy—typically targeting other triatomines, occasionally cockroaches; and iv) coprophagy, consumption of conspecifics’ feces [71–74]. These phenomena enhance the circulation of trypanosomatids, such as dixenous *T. cruzi*, *T. rangeli*, and monoxenous *Blastocrithidia triatomae* [75] but contribute little to the diversification of potential donors. Recent studies have suggested a limited role for sugary food in triatomine nutrition. This includes experimental evidence of sugar ingestion under laboratory conditions—albeit with elevated mortality [76]—as well as the detection of plant DNA in the gut and an amylase gene in the genome of one species [77].

Considering decades of research, the recent emergence of such evidence—primarily from artificial settings or molecular methods—reinforces the conclusion that blood is the sole essential nutrient for these trypanosomatids, with any deviations representing rare, starvation-driven survival strategies.

Thus, the most intensively studied trypanosomes (American *T. cruzi* and *T. rangeli*, and the African salivarian species) are unlikely hosts for RNA viruses, because of the predominantly hematophagous behavior of their vectors. Regrettably, trypanosomes that may be more promising for viral discovery—due to their transmission by vectors with more diverse diets—have been effectively neglected. Although the current knowledge of vectors remains fragmentary, we suggest prioritizing research on species transmitted by Nematocera. In addition to sand flies, this taxon includes four Culicomorpha families with hematophagous representatives: Culicidae (mosquitoes), Corethrellidae (frog-biting midges), Ceratopogonidae (biting midges), and Simuliidae (black flies). These insects exhibit adult feeding habits similar to those in sand flies, while their larvae are (semi)aquatic and either detritivorous or predatory [78, 79]. The trypanosome species transmitted by these vectors belong to the following subgenera: anuran *Trypanosoma*; avian *Trypanomorpha*, *Ornithotrypanum*, and *Avitrypanum*; squamate/mammalian *Squamatrypanum*; as well as mammalian *Megatrypanum* [80–85].

The latter subgenus is of particular interest, as it includes the still unresolved *T. theileri* species complex represented by ruminant-parasitic flagellates. Although usually considered non-pathogenic, *T. theileri*-like trypanosomes have been occasionally associated with severe conditions in cattle (reviewed in [86]). By analogy with LRVs in *Leishmania* spp., yet-undetected RNA viruses could contribute to increased virulence of these trypanosomes. Another promising group is the subgenus *Herpetosoma* vectored by fleas [87], which may also harbor monoxenous trypanosomatids of the genera *Blechomonas*, *Herpetomonas*, and *Leptomonas* [88], in some of which viruses have already been recorded [21]. Expanding viral screening across this diverse range of trypanosomes will not only deepen our understanding of parasite biology, but also shed light on the evolutionary and ecological dynamics shaping virus–host interactions in these protists.

## Conclusions

The discovery of three novel RNA viruses in a single strain of *Trypanosoma platydactyli*, after a long-standing absence of reports from trypanosomes highlights an underexplored aspect of their biology.

The presence of viruses in these flagellates appears linked to vector feeding behavior, which determines the range of potential viral donors. Intensively studied human- and livestock-infecting trypanosomes, vectored by tsetse flies or kissing bugs, are likely poor hosts for RNA viruses. Broader screening of trypanosomes transmitted by Nematocera is expected to significantly expand known viral diversity in trypanosomatids and highlight its ecological and evolutionary drivers.

The discovery of RNA viruses in *Trypanosoma* sheds new light on the evolution of associations between viruses and eukaryotic microbes. Our findings suggest that distribution of viruses is determined primarily by ecological constraints rather than intrinsic cellular barriers. This concept extends beyond trypanosomatids, providing a general framework for understanding the conditions that permit microbial host switching by viruses.

## Methods

### Cultivation, light and electron microscopy

*Trypanosoma platydactyli* strain RI-340 reported in a previous study [27] had been isolated in 2021 from the common wall gecko, *Tarentola mauritanica,* in Bari, Italy (41°03′04″N, 16°53′39″E). Cultivation was performed at 23 °C on biphasic blood agar with an overlay of Schneider’s *Drosophila* medium (Merck/ MilliporeSigma, St. Louis, USA), supplemented with 10% fetal bovine serum (Thermo Fisher Scientific, Waltham, USA), 100 μg/ml streptomycin, and 100 U/ml penicillin (Merck, Rahway, USA).

For morphological studies, blood of an infected gecko and the log-phase trypanosome culture established from it were smeared on glass slides, air-dried, fixed with methanol, and stained with Giemsa. The smears were examined under a light microscope with 100× oil immersion objective and photographed. Trypanosomes were measured on the acquired images using Fiji v. 1.54p [89]. High-pressure freezing transmission electron microscopy of cells in culture followed a protocol described earlier [90].

### Extraction, visualization, and sequencing of dsRNA

Total RNA was extracted with TRIzol (Molecular Research Center, Cincinnati, USA) from approximately 10^9^ cells of the RI-340 strain as well as LRV-positive *Leishmania* (*Viannia*) *guyanensis* MHOM/BR75/M4147 and virus-negative *L. hoogstraali* RHEM/SD/1963/NG-26 (LV31), used as positive and negative controls, respectively [5, 14]. For screening purposes, 50 μg of total RNA were treated with S1 nuclease (Merck/ MilliporeSigma) and DNase I (Thermo Fisher Scientific) followed by LiCl precipitation as described previously [91] to remove single-stranded RNA and residual DNA. The obtained preparations were analyzed in a 0.8% agarose gel post-stained with Midori green (Nippon Genetics Europe, Düren, Germany).

Preparation of dsRNA of the RI-340 strain for next-generation sequencing (NGS) was performed as specified before [23]. Sequencing was performed at Macrogen Europe (Amsterdam, Netherlands) on an Illumina NovaSeq 6000 platform using a TruSeq Stranded Total RNA library (with cytoplasmic and mitochondrial rRNA depletion), generating 10.8 million paired 150 bp reads.

### Sequence data processing

The sequence reads trimmed with Trimmomatic v. 0.40 [92] were assembled with Trinity v. 2.15.2 [93], and mapped back to the resulting assembly with Bowtie2 v. 2.5.4 [94] and SAMtools v. 1.20 [95]. The abundance of RNA fragments, expressed in reads per kilobase per million (RPKM), was estimated with BEDTools v. 2.31.1 [96]. Contigs were identified with BLASTn (BLAST+ v. 2.13.0 [97] against the published genome assembly of *T. platydactyli* (GCA_030849675.1) and BLASTx/DIAMOND v. 2.1.10 [98] against the clustered protein database UniClust50 [99]. Secondary structures were predicted using the IPknot++ v. 2.2.1 web server (Sato et al., 2011). Programmed ribosomal frameshifts were searched for with the PRFect tool [33].

### Phylogenetic analyses

The phylogeny of narnaviruses was inferred by combining the new sequences with those used in a previous study [23], retaining only the single external taxon, *Botourmiaviridae*, as an outgroup. For the reconstruction of LRV phylogeny, sequences of *Pseudototiviridae* were compiled from GenBank matching the set used in an earlier inference [14] and supplemented with sequences of *Phytophthora palustris* toti-like virus, which appears to represent an undescribed lineage within the family [100].

Amino acid alignments for individual genes (RdRp for narnaviruses; capsid and RdRp for LRVs) were generated using MAFFT v. 7.490 [101] and optimized in two steps. The best parameters selected based on the highest ultrafast bootstrap values (1,000 replicates) for basal nodes, as estimated by IQ-TREE v. 2.1.2 [102] using substitution models selected by ModelFinder [103]. In the first step, tested parameters included: i) alignment algorithms L-INS-i, E-INS-i or G-INS-i; ii) BLOSUM matrices with similarity levels of 62, 45 or 35; and iii) gap-extension penalties of 0.123, 0.2, 0.4, 0.6, 0.8, or 1. Before the analysis, the resulting 54 alignments were trimmed with trimAl v. 1.4 [104] with a 0.35 gap threshold. In the second step, the best alignment was trimmed using trimAl with gap and similarity thresholds set to 0.6 and 0.01, respectively, while varying the minimum length of the resulting alignment. This parameter, defined as a fraction of original length, was incremented by 0.1 from 0.1 to 1, followed by 0.02 increments within ±0.1 of the best previous value. The optimality was tested in IQ-TREE as in the first step. The selected parameters for each alignment are shown in Additional file 1: Table S4.

The best final alignment of RdRp of *Narnaviridae* and concatenated alignments of capsid and RdRp of *Pseudototiviridae* were used to infer phylogeny using i) maximum likelihood in IQ-TREE automatically selected substitution model and 1,000 standard bootstrap replicates; and ii) Bayesian approach in MrBayes v. 3.2.6 [105] with the same models, 1,000,000 generations and other parameters set by default. No partitioning was applied, as preliminary testing in IQ-TREE indicated that this approach yielded higher average ultrafast bootstrap values than edge-linked, edge-proportional, and edge-unlinked partitioning schemes.

## Abbreviations

dsRNA: Double-strandedRNA
LRV: Leishmania RNA virus (*Leishmaniavirus*)
ORF: open reading frame
RdRp: RNA-dependent RNA polymerase

## Supplementary Information

**Additional file 1: Table S1.** Alignment parameters and substitution models used in phylogenetic analyses.

**Additional file 1: Table S2.** Measurements of the bloodstream forms of *Trypanosoma platydactyli*.

**Additional file 1: Table S3.** Measurements of cells in culture.

**Additional file 1: Table S4.** Information on the viral genomes.

**Additional file 2: Fig. S1.** Full-size image of the gel shown in Fig. 3A, with the cropping area indicated by a dashed outline.

**Additional file 3: Fig. S2.** The region of overlap between capsid and RdRp ORFs in LRVs from *T. platydactyli* and *Leishmania aethiopica*.

## Declarations

### Ethics approval and consent to participate

Not applicable

### Consent for publication

Not applicable

### Data availability

New sequences reported here are available from the GenBank under accession numbers PV052907–PV052909. All other data generated or analyzed during this study are included in this published article and its supplementary information files.

## Acknowledgements

The authors thank members of their laboratories for stimulating discussions.

## Authors’ Contributions

VY conceived and supervised the study; AYK, JAMR, and JV analyzed morphology; DK and JV conducted experiments; AYK and DG performed bioinformatic analyses and interpreted results. AYK, DG, JAMR, and JV prepared illustrations. AYK wrote the manuscript, which was then edited by JL, JV and VY. All authors read and approved the final manuscript.

