## Supplementary figures and images for "The first RNA viruses detected in a trypanosome: novel narnaviruses and a leishmaniavirus in the gecko parasite *Trypanosoma platydactyli*"

### Additional file 2: Figure S1

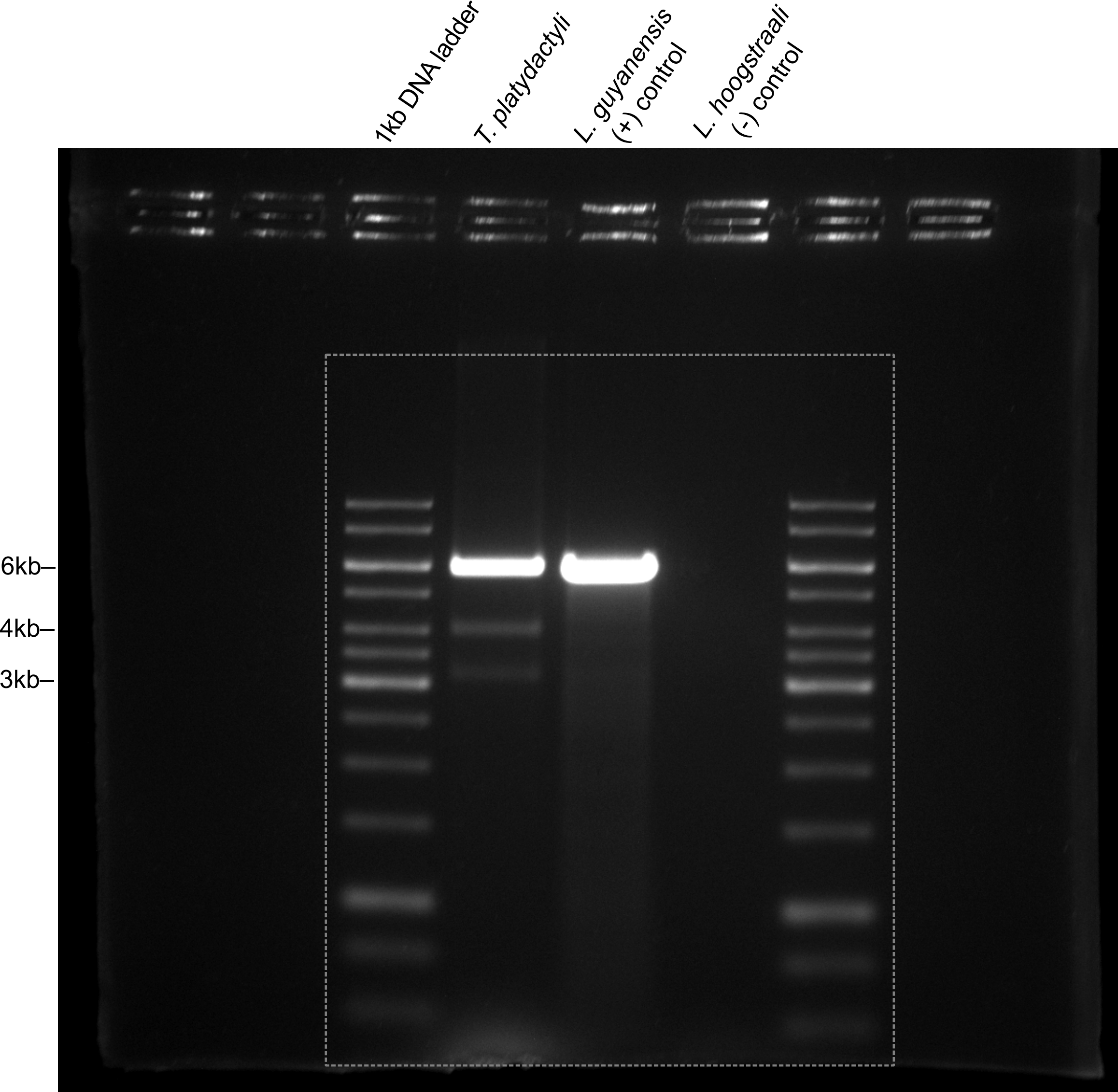

### Additional file 3: Figure S2

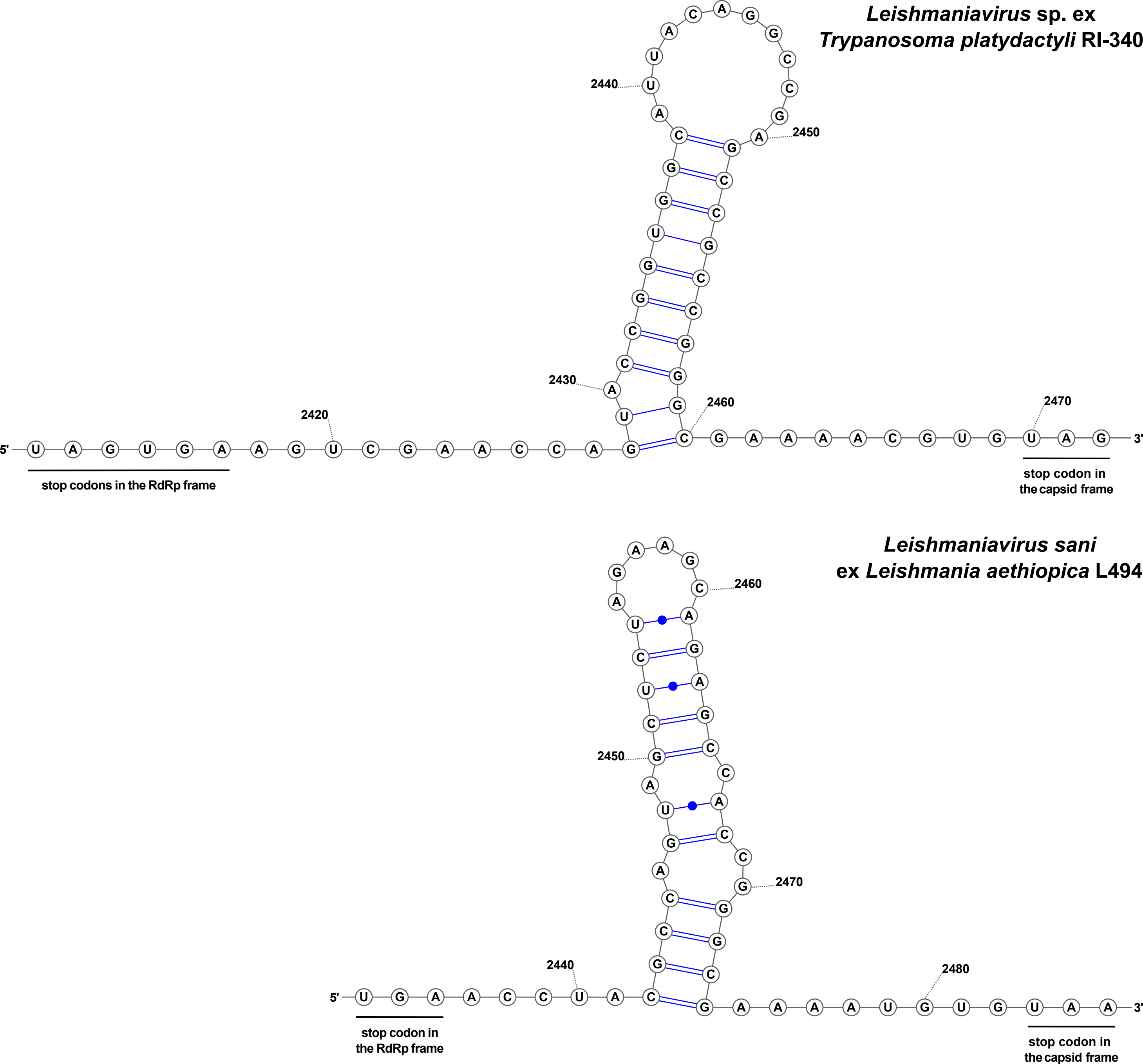
